# Zebrafish Avatar-test predicts patient’s tumor response to chemotherapy in breast cancer: a co-clinical study towards personalized medicine

**DOI:** 10.1101/2024.10.03.616349

**Authors:** Raquel V. Mendes, Joana M. Ribeiro, Helena Gouveia, Cátia Rebelo de Almeida, Mireia Castillo-Martin, Maria José Brito, Rita Canas-Marques, Eva Batista, Celeste Alves, Berta Sousa, Pedro Gouveia, Miguel Godinho Ferreira, Maria João Cardoso, Fatima Cardoso, Rita Fior

## Abstract

Chemotherapy remains the mainstay in most high-risk breast cancer (BC) settings, with several equivalent options of treatment. However, the efficacy of each treatment varies between patients and there is currently no test to determine which option will be the most effective for each individual patient. Here, we developed a fast in-vivo test for BC therapy screening: the zebrafish patient derived xenograft model (zAvatars), where in-vivo results can be obtained in just 10 days. To determine the predictive value of the BC zAvatars we performed a clinical study, where zAvatars were treated with the same therapy as the donor-patient and their response to therapy was compared. Our data shows a 100% correlation between patient’s clinical response to treatment and its matching zAvatar. Altogether, our results suggest that the zAvatar model constitutes a promising in-vivo assay to optimize cancer treatments in truly personalized manner.

## 1. Introduction

Breast cancer (BC) exhibits high heterogeneity at clinical, histological, and molecular levels.^[1–4]^ Depending on the receptor status (ER, PR, HER2), tumors are classified into Luminal A-like, Luminal B-like HER2-, Luminal B-like HER2+, HER2-type, and Triple Negative (TNBC) subtypes.^[1,5,6]^ This classification, together with clinicopathological variables such as tumor stage and grade, has facilitated the development of targeted treatments, leading to significant improvements in outcomes.^[3,7]^

Nevertheless, chemotherapy remains the primary treatment approach in the vast majority of high-risk early BC, and also an important option for metastatic disease. For the treatment of HER2 negative metastatic BC, there are a few standard therapies, mainly for 1^st^ and 2^nd^ line. However, most subsequent treatment options have similar effectiveness and are administered sequentially. The choice of treatment is often based on factors such as the physician’s experience, toxicity profiles, and patient preferences, rather than the sensitivity of the patient’s tumor cells to each specific treatment. This strategy potentially exposes patients to unnecessary toxicities, especially in cases without tumor response, and when other therapeutical options are available.

Genomic profiling and biomarker development have been the hope for personalized and precision cancer treatment.^[8]^ However, the recent recognition of genetic and non-genetic cancer heterogeneity, particularly in BC,^[7,9,10]^ is a significant stumbling block for a personalized approach. Therefore, there is a clear unmet need for a direct functional sensitivity test i.e., challenge directly the tumor cells with the available therapeutical options to determine the most cytotoxic treatments. Mouse patient-derived xenografts (mAvatars) have emerged as essential tools in translational cancer research to predict tumor response to treatment.^[11–13]^ However, establishing mAvatars and conducting compound testing can take months, making it impractical for clinical advice.^[14]^ Alternatively, in-vitro cancer organoids and spheroids/whole-tumor cell cultures are emerging as promising models.^[15–18]^ Still, these models do not have the complexity of a living organism, limiting the test of several therapies (such as antiangiogenic, or therapies that need *in vivo* metabolization) and more important the evaluation of the tumor metastatic potential.^[19–22]^

Our strategy was to develop an intermediate approach: a rapid in-vivo assay with unparalleled cellular resolution – the zebrafish patient-derived xenograft model or zAvatar-test.^[19,22,23]^ This assay relies on the injection of fluorescently labelled tumor cells into zebrafish and access tumor behavior and response to therapy after 2-4 days of treatment. The overall test including single-cell confocal microscopy imaging analysis takes 10 days. Zebrafish xenografts provide speed, cellular resolution, and the capacity to perform numerous transplants. They also enable the assessment of critical cancer hallmarks, including metastatic and angiogenic potentials, thanks to the significant genetic similarity between humans and zebrafish.^[19,22–24]^

In the present study, we challenged zAvatars to discriminate different cancer sensitivities to BC therapies and evaluate the predictive power of the model to forecast patient’s response to therapy. To optimize the model, negative and positive controls were first generated with TNBC cell lines to screen the major therapeutic options present in the most commonly used international guidelines.^[6,25,26]^ Next, we performed a clinical study where patient-derived BC zAvatars were treated with the same therapy as the patient, to compare treatment response of each patient to their matching zAvatar-test.

## 2. Results

### 2.1 TNBC zebrafish xenografts respond differently to Anthracyclines and Taxanes

In order to develop the BC zebrafish xenograft model, we started by selecting two different TNBC cell lines that have been extensively characterized in-vitro and in-vivo.^[19,27–29]^ Our first goal was to establish these cancer models and screen the major chemo-therapeutic options present in the international guidelines, generating positive and negative controls for each therapy, to then focus on patient samples.

Consistent with previous reports,^[19,27–29]^ MDA-MB-468 with grape-like structure present a higher percentage of Ki-67 and mitotic index than the stellate Hs578T tumors (Figure. 1A and Figure. S1A). Hs578T showed a higher level of basal apoptosis (activated Caspase3) in relation to MDA-MB-468 (Figure 1A and Figure S1A). Both TNBC xenografts presented micrometastasis in multiple organs of the zebrafish embryo (Figure 1B and Figure S1A).

**Figure 1.**
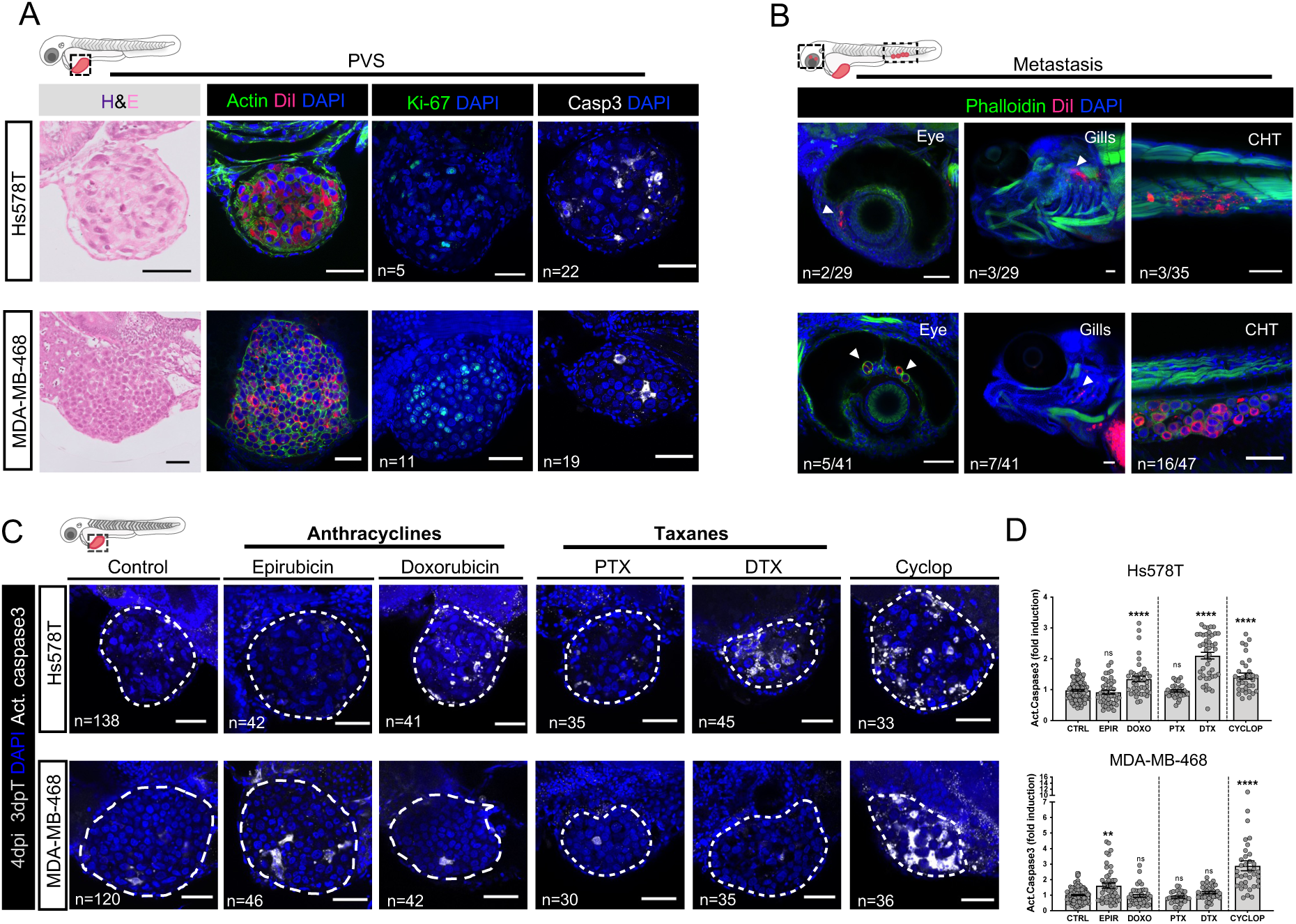
Characterization and comparison of drug sensitivities in human BC tumor zebrafish xenografts. (A) Characterization of TNBC zebrafish xenografts. Hs578T and MDA-MB-468 cancer cells were labelled with DiI (red) and injected into the perivitelline space (PVS) of 2 days post-fertilization (dpf) zebrafish embryo. 4 days post-injection (dpi) zebrafish xenografts were analyzed through H&E and whole-mount immunofluorescence for Actin, Ki-67 and apoptosis (activated caspase3) (white). (B) At 4dpi tumor cells can be detected at distant sites, forming micrometastasis in the eyes, gills (arrowheads) or caudal hematopoietic tissue (CHT). (C) 2dpi zebrafish xenografts were treated in-vivo with epirubicin, doxorubicin, paclitaxel (PTX), docetaxel (DTX), or cyclophosphamide and compared with untreated controls. (D) Apoptosis (activated caspase3 in white) was analyzed and quantified (epirubicin: FI apoptosis Hs578T=0.82 vs MDA-MB-468=1.33; doxorubicin: FI apoptosis Hs578T=1.24 vs MDA-MB-468=0.83; PTX: FI apoptosis Hs578T=0.90 vs MDA-MB-468=0.83; DTX: FI apoptosis Hs578T=2.13 vs MDA-MB-468=1.1; Cyclop: FI apoptosis Hs578T=1.32 vs MDA-MB-468=2.43) (\**\*P=* 0.0018). Data represents AVG±SEM and are averages from 3 independent experiments. Statistical analysis was performed using Mann–Whitney test. Statistical results: (ns) > 0.05, \*\**P* < 0.01, \*\*\*\**P* < 0.0001. White dashed line delimits the tumors. Total number of analyzed xenografts is indicated in the images. Scale bars:100μm (H&E) and 50μm (immunofluorescent images). All images: anterior to the left, posterior to right, dorsal up and ventral down.

The standard of care for breast cancer neoadjuvant chemotherapy comprises the sequential administration of an anthracycline (epirubicin or doxorubicin) with cyclophosphamide, followed by a taxane (docetaxel or paclitaxel).^[30]^ The 2 anthracyclines and the 2 taxanes are considered equivalent compounds,^[6]^ since in several studies the average responses were similar between these compounds of the same family.^[31,32]^

First, we started by evaluating the responses to the single agents and then to the combinations (see methods and Figure S2). To achieve this, at 1-day post injection (1dpi), TNBC xenografts were randomly distributed between control-untreated and treatment groups. To determine the anti-tumor efficacy of the compounds, xenografts were processed for immunofluorescence and imaged by single cell confocal microscopy to determine cell death by apoptosis (activated caspase3) (Figure 1A and Figures S2B and S2C).

Surprisingly, our data reveal that TNBC xenografts can respond in opposing ways to the two equivalent anthracyclines treatments. Hs578T tumors were resistant to epirubicin but sensitive to doxorubicin, whereas MDA-MB-468 xenografts were only sensitive to epirubicin (Figure 1C and 1D).

Regarding the taxanes, both TNBC models were resistant to paclitaxel (PTX) whereas docetaxel (DTX) induced a significant increase in cell death in Hs578T but not in MDA-MB-468 xenografts (Figure 1C and 1D). Of note, colorectal SW620 xenografts were sensitive to PTX (Figure S1B and S1C) demonstrating that the lack of response in the TNBC cells was not a technical issue of the model. These results show that it is possible to distinguish sensitivities to different anthracyclines and taxanes.

In addition, we also tested Cyclophosphamide (Cyclop) as a single agent (Figure 1C and 1D) and in combination with anthracyclines followed by taxanes (Figure S2A - S2C). Cyclophosphamide alone induced a massive cell death in both TNBC models (Figure 1B and 1D), and this was further enhanced when combined with the other compounds (Figure S2B and S2C). These results are consistent with numerous preclinical and clinical studies, showing that the use of combination chemotherapy significantly improves clinical and pathologic response rates rather than single agents.^[33]^ Overall, our data demonstrate the capacity of the model to distinguish tumor responses, even to compounds belonging to the same drug family.

### 2.2 BC zAvatars can be efficiently established from early and advanced BC tumors and preserve relevant BC biomarkers

After establishing the assay, we moved to optimize BC zebrafish patient-derived xenografts (zAvatars). We used BC surgical samples, core needle biopsies and biological fluids (ascites, malignant pleural effusion) from adenocarcinomas of diverse tumor stages and subtypes (Luminal, HER2 enriched and TNBC) (Table S1 and S2). Tumor cell suspensions were injected into 2days post-fertilization (dpf) zebrafish embryos without prior in-vitro expansion. We successfully implanted 51 out of 52 patient samples into the zebrafish embryos from early (n=40) and from advanced (n=12) stages. zAvatars displayed very heterogenous implantation rates, (Figure 2A and Table S2); being implantation defined as the frequency of zebrafish xenografts that maintain a tumor mass at the end of the assay (3-5 dpi).

**Figure 2.**
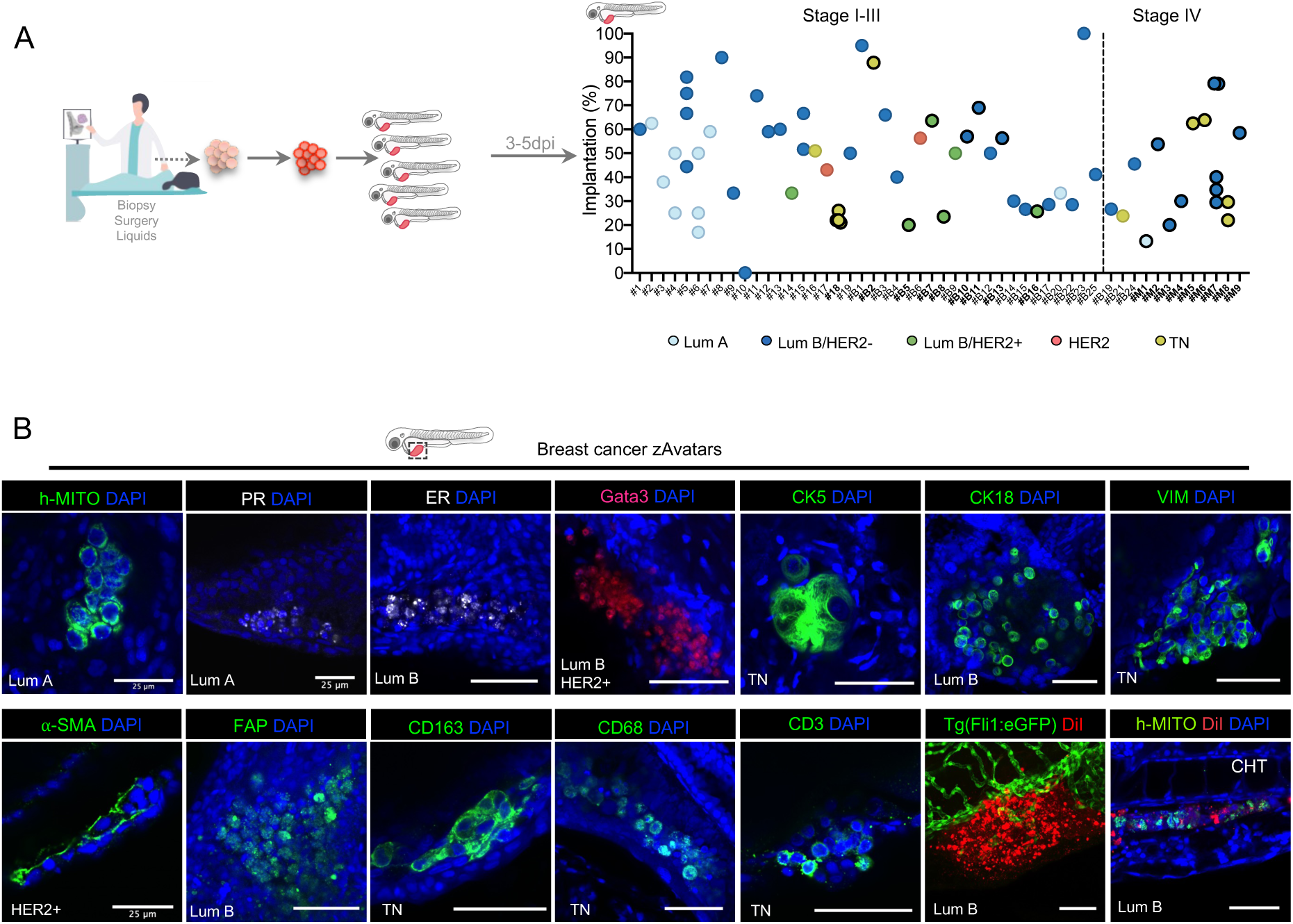
Characterization of BC zAvatars. (A) Workflow summarizing the zAvatar assay. Implantation rate at 3, 4 or 5dpi and the corresponding receptor status of the tumors from breast core needle biopsies, surgical samples, and liquids. (B) zAvatars were processed for whole-mount immunofluorescence to assess human-associated antigens (anti-human mitochondria), and BC biomarkers: PR, ER, Gata3, CK5, CK18, mesenchymal marker: VIM, stromal components: a-SMA, FAP, immune cells: CD163, CD68, CD3, angiogenesis and micrometastasis potential. ER-Estrogen Receptor, PR-Progesterone receptor, Vim-Vimentin, FAP-Fibroblast activation protein, a-SMA-alpha smooth muscle actin, CK-Cytokeratin, CHT-Caudal hematopoietic tissue. # - surgical sample, #B - breast biopsy, #M – Metastatic (either biopsy of metastasis or liquids from metastatic patients). Darker circles represent samples used for matching patient’s response to therapy. Scale bars:50μm, except if identified has otherwise. All images: anterior to the left, posterior to right, dorsal up, and ventral down.

We also detected the expression of various BC and tumor microenvironment (TME) markers, in the different cancer subtypes (Figure 2B), demonstrating that zAvatars faithfully conserve the most relevant BC-associated biomarkers, even the elusive ER. Overall, we successfully established BC zAvatars derived from all 5 different BC subtypes and from various types and stages of BC, including treatment-naïve, drug-treated, primary, metastatic, biopsies and surgical specimens and biological fluids.

### 2.3 BC zAvatars can predict tumor response to treatment

Next, we designed a clinical study to evaluate the predictive value of the BC zAvatar-test. In this study, we treated the zAvatars with the same therapy as the donor-patient and compared the patient clinical response with its matching zAvatar (Figure 3A). We generated zAvatars from 18 patients, ranging from early (stage II-III) (8 from core needle breast biopsies and 1 from a surgical specimen) to advanced BC (stage IV) (6 core needle biopsies, 2 pleural effusions and 1 ascites) (Figure S3A).

**Figure 3.**
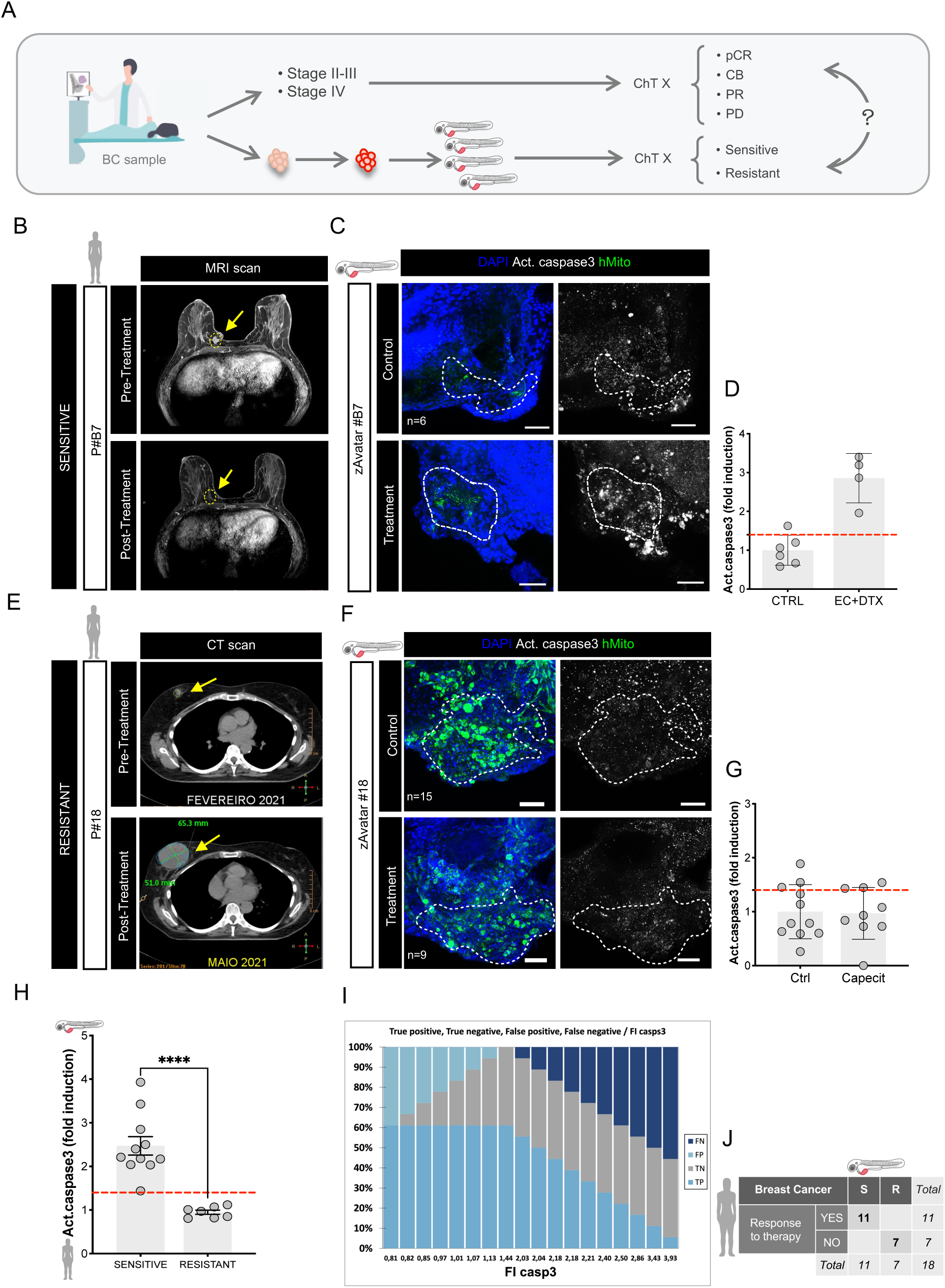
zAvatars from early and advanced BC can predict patient response to treatment. (A) Workflow of the clinical study. Patients’ samples were injected into zebrafish embryos, and zAvatars were subsequently treated with the same chemotherapy regimen as their respective donor-patients. Treatment responses of the zAvatars were then compared to the clinical response of their corresponding patients. (B) Patient MRI (P#B7) before and after treatment. (C) Corresponding BC zAvatars from core needle-biopsy (#B7), treated in-vivo with the same ChT regimen as its donor-patient and compared with untreated controls. 3dpi zAvatars, corresponding to 2 days post-treatment (dpt). (D) Apoptosis (activated caspase3) was analyzed and quantified. (E) Patient CT scans (P#18) before and after treatment. (F) Corresponding zAvatars from surgery sample (#18) treated in-vivo with the same ChT regimen as its donor-patient and compared with untreated controls. zAvatars fixed at 3dpi, corresponding to 2 dpt. (G) Apoptosis (activated caspase3) was analyzed and quantified. (H) Activated Caspase3 fold induction of zAvatars and its corresponding matching patient’s response segregated according to sensitive (pCR, PR, CB) and resistant (PD). (I) ROC curve graph (J) Confusion matrix displaying the number of patients that did responded (YES) or did not respond (NO) to the treatment, along with the corresponding zAvatar-test: sensitive (S) or resistant (R). Data represents AVG±SD and are averages from one independent experiment. Each dot represents one xenograft. Red dashed line represents the threshold value of activated caspase3 induction calculated by ROC curve analysis. Statistical analysis was performed using Mann–Whitney test. Statistical results: (ns) > 0.05, \*\**P* < 0.01. pCR - pathologic complete response; PR - partial response; PD - progressive disease; CB - clinical benefit; PD – progressive disease. CT – computerized tomography; MRI – magnetic resonance imaging. Apoptosis (activated caspase3 in white, Z-projection image). Yellow arrows points to the tumors. Scale bars:50μm. All images: anterior to the left, posterior to right, dorsal up, and ventral down.

Clinical response to treatment was determined based on clinical assessment, imaging, and histological confirmation, when appropriate (see Methods). Patients whose tumors achieved a pathological complete response (pCR) or a partial response (PR) in the early setting, or a clinical benefit (CB=complete/partial response and stable disease) in the advanced setting were considered sensitive to treatment; patients whose tumors presented progressive disease (PD) were considered resistant to treatment. All zAvatar-test data obtained was blindly compared with the patient clinical outcome (see methods).

Patient tumor cells were processed without in-vitro expansion and generated zAvatars were randomly distributed into 2 groups: control (untreated) and treatment (same chemotherapy as the respective patient). zAvatars were processed for immunofluorescence and induction of apoptosis (activated caspase3) after treatment was quantified to define sensitivity or resistance to treatment.

In Figure 3 we show 2 examples of patients with different treatments outcomes and their corresponding zAvatar-test results. Patient #B7, initially diagnosed as cT1c(m)N1cM0, underwent neoadjuvant combination therapy with EC+Docetaxel. Following treatment completion, the tumor achieved a pCR (Figure 3B). The corresponding zAvatars test, showed 2.85-fold induction of apoptosis (activated caspase3) upon treatment when compared to untreated zAvatars controls (Figure 3C, D and Figure S3B).

In contrast, patient#18, staged as ypT3N0M0 at the surgery, received adjuvant Capecitabine treatment but later the disease progressed with lung metastasis (Figure 3E). The zAvatar-test showed a lack of response to the treatment (Figure 3F, G and Figure S3B).

zAvatars from patients whose tumors were sensitive to treatment showed a statistically significant higher average Apoptosis Fold Induction (AFI) than zAvatars from patients whose tumors progressed (Figure 3H). These findings highlight a robust correlation between AFI in zAvatars and tumors’ response to treatment, regardless of tumor stage.

To evaluate the capacity of zAvatars to differentiate between clinical sensitivity and resistance in a binary manner, we conducted a Receiver Operating Characteristic (ROC) analysis (Figure 3I, Figure S3D and S3E). This analysis was based on the average apoptosis AFI observed in zAvatars (Figure 3D, 3G, S3B and S3C). The ROC analysis revealed that the optimal threshold for distinguishing sensitivity and resistance in zAvatars was 1.44-FI of apoptosis, with the highest specificity and sensitivity performance (AUC = 1). Based on this threshold of apoptosis, all 18 zAvatars tests matched their donor-tumor response to treatment. The zAvatars with an AFI equal or above 1.44 were categorized as sensitive (S) to therapy, and their corresponding donors achieved a pCR, PR, or CB as a treatment outcome. Conversely, the zAvatars with a AFI below 1.44 were classified as resistant (R) to therapy, and their donors did not respond to the treatment (PD) based on their clinical evaluation (Figure S3B and S3C). Considering the defined cut-off value, Figure 3J presents a confusion matrix displaying the number of patients with actual vs predicted responses in zAvatars. These findings indicate that the zAvatar-test accurately predicted the treatment response of its corresponding donors with a 100% accuracy rate, in all 18 patients.

As a limitation, the zAvatar model was insufficient to discriminate between pCR and PR. Partial responses can be very heterogenous, exhibiting a wide range of tumor reduction, ranging from 36.1% (#B10) to 92.5% (#B16) (Figure S3B and Table S2), and their corresponding zAvatar-test was sensitive, not being able to discriminate these levels of response.

Altogether, these results show that the zAvatar-test was able to predict its donor-patient treatment outcome (sensitive/resistant to treatment) in 100% of cases in early and advanced stages of the disease (Figure 3J, Figure S4A and S4B).

### 2.4 zAvatar micrometastasis potential correlates with patient’s original tumor stage

One of the major advantages of the zAvatar model is the possibility to evaluate cell dissemination and micrometastasis in-vivo (Figure 4A). Thus, we next evaluated whether zAvatars were able to reflect the metastatic potential of its donor-patient tumor sample (early or metastatic). We analyzed not only the previous cases used for therapy correspondence, but also patients that had been recruited to the study but were not included due to several reasons (e.g., no treatment or changed hospital) (Figure S5A).

**Figure 4.**
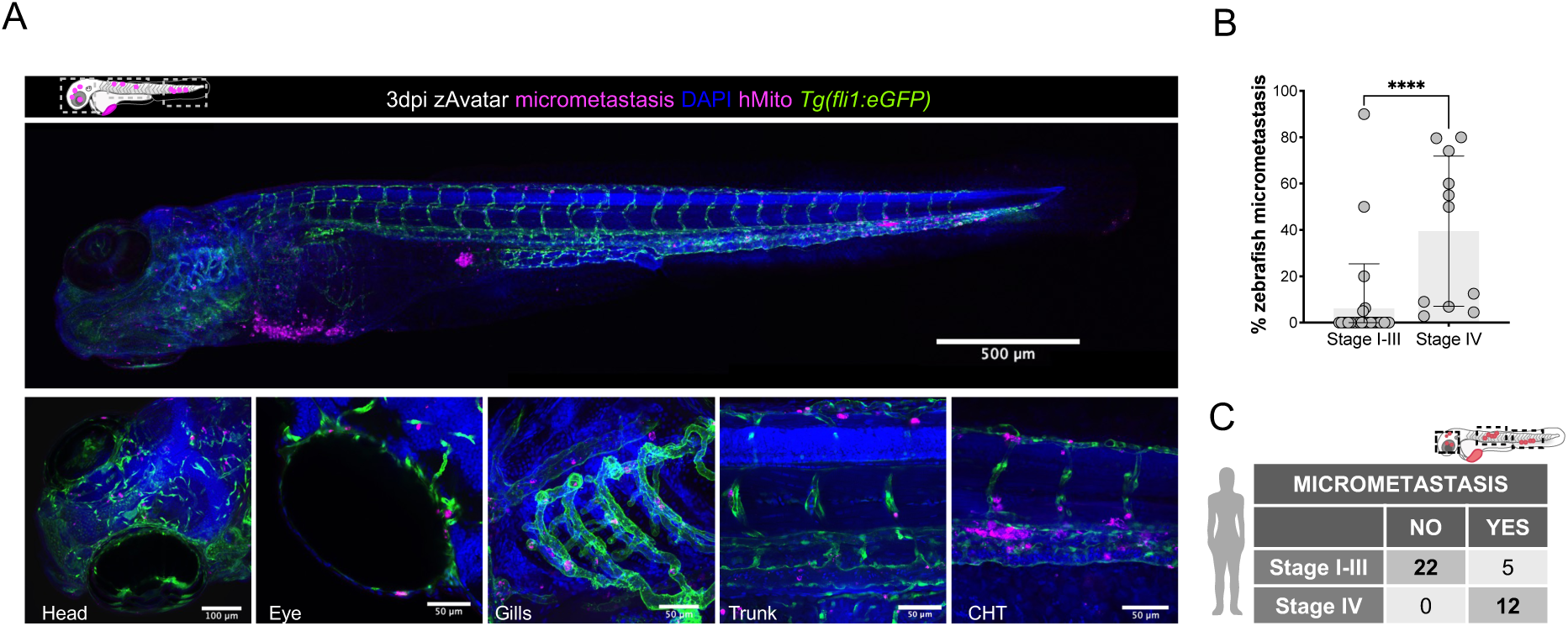
zAvatars from early and late-stage BC can recapitulate its donor-patient tumor origin. (A) 3dpi zebrafish Avatar showing tumor cells. (B) Incidence of micrometastasis in untreated zAvatars at 3-5dpi derived from early and late-stage patients (\*\*\*\**P*<0.0001). Results are from one independent experiment. Each dot represents one xenograft and its corresponding number of disseminated cells. Bars are AVG±SD. (C) Confusion matrix showing that metastatic potential correlates with tumor staging. Statistical analysis was performed using Mann– Whitney test. CHT-caudal hematopoietic tissue. All images: anterior to the left, posterior to right, dorsal up, and ventral down.

Among the zAvatars derived from early BC stages (local stage I-III), micrometastasis was not detected in most cases (22/27; 81.48%) (Figure 4B, C and Figure S5A). In contrast, all zAvatars derived from metastatic BC (stage IV) exhibited cell dissemination and formation of small micrometastasis (12/12; 100%) (Figure 4B, C and Figure S5A). These micrometastasis varied from a few cells to dozens or even hundreds (Figure S5A). These findings indicate that, in general, the metastatic potential of zAvatars matches with the corresponding tumor stage of its donor-patient (Figure 4C and Figure S5A).

In most BC cases, it can take years or even decades from the initial diagnosis for metastasis to become clinically evident.^[34,35]^ Nevertheless, we questioned if the presence of micrometastasis in the zAvatars could correlate with future tumor progression/metastasis in the patient. For this we analyzed only early-stage patients (I-III) with non-metastatic disease at the time of surgery/sampling. From 27 zAvatars analyzed, 22 of them did not show micrometastasis. Their corresponding patients also did not develop metastasis (1-5 years since biopsy; median follow up) (Figure S5A and Table S1). However, in the remaining 5 zAvatars, in 4 of them (#B5, #B12, #B13 and #B20), we could identify single cell dissemination and in 1 case (#18) several micrometastasis. These 4 patients (stage II (#B12 and #B20) and III (#B5 and #B13)) underwent neoadjuvant therapy, resulting in partial or complete response with tumor reductions ranging from 70% to 98%. These 4 patients are currently metastasis-free (median follow-up of ∼51 months). In contrast, in the case of zAvatar #18, we observed the presence of several micrometastasis. The donor-patient, diagnosed with stage II TNBC, underwent neoadjuvant systemic therapy, and the tumor sampling for zAvatars was conducted during surgery. The patient presented a significant residual disease burden (ypT3N0M0; RCB score 2) after treatment, which is known to be associated with a higher risk of metastatic recurrence.^[36]^ Metastatic progression in the lung was objectivized 5 months after surgery under adjuvant systemic treatment.

It is still premature to evaluate whether the zAvatar model can predict which patients will develop metastasis in the future, since cells can remain dormant for decades in patients, requiring a longer follow-up to observe potential metastatic events that may occur.

Nevertheless, these findings suggest that zAvatars have the potential to enhance risk stratification in BC patients following neoadjuvant systemic treatment. With the current range of treatment options available in the post-neoadjuvant setting, it is conceivable that personalized response evaluation using zAvatars may contribute to informed treatment decisions and ultimately improve patient outcomes. This highlights the promising role of zAvatars in optimizing therapeutic strategies for BC patients.

## 3. Discussion

Despite recent advances, BC treatment response varies widely due to heterogeneity in cell and molecular characteristics of BC tumors, even within the same histological subtype.^[37,38]^ A test that could evaluate individual’s tumor response to the therapeutic options prior to treatment initiation, is unquestionably an unmet-need in cancer treatment.

Here we developed and validated the BC zAvatar model for personalized medicine.

Although BC is considered the most difficult type of cancer for tumor establishment in mouse models,^[11,39–41]^ we were able to generate BC zAvatars without in-vitro expansion. We successfully established zAvatars from BC patients representing all 5 different subtypes, stages of the disease and independently on their aggressiveness. zAvatars from ER positive tumors were successfully generated and these maintain the ER expression, which can be challenging in mAvatars.^[12,13,18,40,42–48]^ Moreover, we shown that zAvatars can retain specific BC biomarkers like ER, PR and Gata3, and preserve their TME.

More importantly, we showed that zAvatars accurately predicted the donor response to treatment in all 18 cases. This was observed across various therapeutic options (either single or combinations) and disease stages, demonstrating the versatility of the zAvatar-test.

Finally, we showed that the majority of zAvatars from patients with early BC did not form micrometastasis, whereas all advanced BC-derived zAvatars presented micrometastasis, suggesting that the metastatic potential in the zAvatars matches the tumor stage. These are still small numbers and therefore it is still very early to evaluate whether the zAvatar model offers the opportunity to predict BC recurrence, since it can take years to decades for the disease to develop metastasis after the original diagnosis. Metastatic risk assessment would be extremely helpful to improve BC treatment management.

Considering the limitations of the zAvatar model, the main limiting factors are availability and size of fresh tumor samples, since we do not amplify tumor cells in culture to avoid artifacts of in-vitro selection and guarantee fast results. The neoadjuvant setting is demanding due to the small amount of tumor material that is taken from a core needle biopsy. This is a challenge, not just for zAvatars but for every functional model;^[46,49]^ nonetheless, the recent whole-tumor-cell culture model may be a good alternative given their fast assay and predictive value.^[15]^

Another limitation is that the zAvatar-test couldn’t distinguish between a pCR and PR. In this study, PRs were very heterogeneous ranging from 36.1% to 98% of tumor reduction in the patient setting. Furthermore, it is important to note that the pathological assessment of neoadjuvant treatment includes evaluation of the response in the lymph nodes. However, we did not have access to the lymph nodes prior to treatment, as we only received biopsies from the primary lesion.

Nevertheless, considering the cut-off value defined by the ROC curve analysis, our zAvatar-test was able to efficiently predict the treatment response in 18 out of 18 BC patients, in early and advanced stage disease. Even though the sample size may seem not very extensive, the total number of zAvatars and matching patients surpasses that of any other published BC model, to our knowledge,^[11–13,15–17]^ (mAvatars (12/13),^[11,13]^ PDO (3/3),^[50]^ and whole-tumor cell cultures (15/15)).^[15]^

Moreover, we must highlight the fact that our assay not only used BC samples from different stages and from different sources, as it also allowed the evaluation of the sensitivity of several different combinatorial treatments, including therapies that can only be metabolized in-vivo, like cyclophosphamide,^[49,51]^ and methotrexate.^[52]^ For these therapeutic agents, organoids, or *ex-vivo* cultures cannot be used as predictors for treatment response, limiting their use for precision therapy.

In summary, we have generated a BC zAvatar-test from early and advanced stages, from different types of samples, encompassing diverse molecular subtypes. Importantly, the zAvatar-test demonstrated a high level of accuracy in mirroring the therapeutic responses observed in patients. These findings provide strong evidence for the potential use of zAvatar-test in guiding personalized treatment for BC patients in the future. Our results also show that the zAvatar metastatic potential correlates with tumor staging. Although still early, further data and time are required to assess the potential impact of the absence of micrometastasis in zAvatars on refining risk stratification in BC. Similarly, the presence of micrometastasis in zAvatars may provide valuable insights into determining which patients should undergo more intensive/targeted adjuvant therapy, as well as a more comprehensive surveillance program.

## 4. Materials and Methods

### Zebrafish care and handling

In-vivo experiments were performed using zebrafish (*Danio rerio*), nacre, casper and Tg(Fli1:eGFP), which were handled according to European animal welfare Legislation, Directive 2010/63/EU (European Commission, 2016).

### Human Breast Cancer cell lines

Human breast cancer cell lines, MDA-MB-468 and Hs578T, were kindly provided by Mónica Bettencourt-Dias laboratory at Instituto Gulbenkian de Ciência, Portugal. All cell lines were authenticated through short tandem repeat profile analysis and tested for mycoplasma.

### Cell Culture

BC cell lines were cultured and expanded to 70-80% confluence in Dulbecco’s Modified Eagle Medium (DMEM) High Glucose (Biowest) supplemented with 10% Fetal Bovine Serum (FBS) (Sigma-Aldrich) and (1%) Penicillin-Streptomycin (P/S) (10,000 U.mL^-1^) (Hyclone). For Hs578T, culture medium was supplemented with insulin at 10μg/mL (Sigma-Aldrich). BC cells were cultured in a humidified atmosphere containing 5% CO^2^ at 37**°**C.

### Cell Labelling

Cells were labelled with the lipophilic dye vybrant CM-DiI (Vybrant^TM^ CM-DiI Thermo Fisher Scientific) at a (2μL.mL^-1^) concentration according to manufacturer’s instructions. BC cells were resuspended to a final concentration of (0.50×10^6^cells.μL^-1^).

### Human Tissue Processing

The study was approved by the Ethics Committee of the Champalimaud Foundation.

Human BC resected samples and biopsies used for zAvatars establishment were obtained from Champalimaud Clinical’s Centre (CCC) Breast Unit, with written informed consents. We based this BC sample preparation protocol on our previous experience in colorectal cancer,^[22]^ and protocols on human BC organoids procedures, that allow to maintain stemness and cell viability during processing.^[17,53]^ Primary tissue from surgical resected breast cancer samples and biopsies were collected (Supplementary Table 2) and cryopreserved in (90% FBS and 10% DMSO). When defrosted, tumor tissues and biopsies were minced in Mix1 (Supplementary Table 2). Subsequently, breast tissue was digested with Collagenase (20mg×mL-1, Worthington) and Hyaluronidase (3mg×mL^-1^, Sigma) for 10 min at 37°C. Tumor cell suspension was filtered and centrifuged at 200xg for 4 min. For cell labelling, tumor cells were incubated with the fluorescent cell tracker DiI (10 μL·mL^−1^) in Mix2 (Supplementary Table 2), for 15 min at 37°C and then for 5 min on ice. Tumor cells were checked for viability with trypan blue dye exclusion. Cancer cells were resuspended in Mix1 with human EGF (5 ng·mL^-1^, Peprotech), FGF7 (5 ng·mL^-1^, Peprotech), FGF10 (20 ng·mL^-1^, Peprotech), A83-01 (500nM, Peprotech) and Neuroregulin-1 (5nM, Peprotech). Same protocol was used for liquids processing (Supplementary Table 3). All cancer cells were resuspended to a final concentration of (∼0.25 × 10^6^ cells mL^-1^) and a small aliquot of the processed/dissociated tumor sample was stained with MGG Grunwald-Giemsa (Bio-Optica) method according to the manufacturer’s instructions.

### Clinical response assessment

Tumor staging was performed according to the American Joint Committee on Cancer (AJCC) Cancer Staging Manual, 8th edition.

### Evaluation of response to treatment in patients with early breast cancer

In patients undergoing neoadjuvant systemic treatment breast and nodal disease were assessed by physical examination, breast imaging, and histopathologic diagnosis with Ki-67, ER, PR and HER2 receptor status assessed on pre-treatment core biopsy. Hormonal receptor and HER2 status was defined according to ASCO/CAP guideline.^[54]^ ER or PR were considered positive if more than one percent of tumor cells stained. Tumors were considered HER2 positive if they were 3+ by immunohistochemistry or demonstrated gene amplification by *in situ* hybridization. A pCR was defined as no invasive residual tumor in breast and axillary lymph nodes removed following neoadjuvant therapy. Partial response was defined as a non-pCR response. Clinical response was assessed by clinicians-investigators independently of the zebrafish Avatar drug sensitivity results. Experimental researchers had no previous information about the clinical outcome of any patient at the time of the experiment, only had information regarding the type of therapy given to the patient. Then, the zAvatar results along with the anonymized patient identification codes were sent to the clinicians-investigators to compare patient clinical response with the zAvatar-test.

### Evaluation of response to treatment in patients with metastatic breast cancer

Response evaluation in the metastatic setting was performed by physical examination, blood tests and imaging exams, usually PET-CT or CT-scans, according to clinical practice: at baseline (before initiating systemic treatment) and every 2 to 4 cycles of treatment. Image was independently assessed for clinical response by a dedicated radiologist or nuclear medicine expert, blinded to drug sensitivity results. Responses were evaluated using the Response Evaluation Criteria in Solid Tumors (RECIST) version 1.1 criteria.

### Zebrafish Xenografts Injection

Labelled cells were injected into the perivitelline space (PVS) of anesthetized 2 days post fertilization (dpf) zebrafish larvae. Following microinjection, all larvae were maintained at 34°C until the end of the experiments in E3 medium. At 1 day post injection (dpi), zebrafish xenografts were screened regarding the presence or absence of a tumoral mass. All xenografts were kept at 34°C until the end of the experiment. At 3, 4 or 5 dpi (depending on the chemotherapy scheme), xenografts were sacrificed and fixed with 4% formaldehyde (Thermo Scientific) and kept in 100% methanol at −20°C for long-term storage. We designed the test to span maximum 5 days not only due to animal ethics constrains but more importantly to give time to perform immunofluorescence, confocal imaging, and analysis in a useful time window for future patient advice. The tumor implantation percentage was calculated as follows:

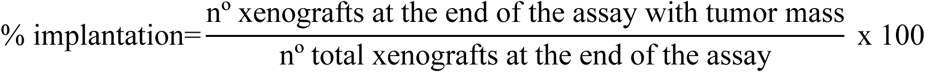

### Zebrafish-Xenograft Drug administration

Zebrafish larvae Maximum Tolerated Concentration (MTC) of each compound (Table S2), was determined using as reference the maximum patient’s plasma concentration as referenced in (Table S2). For Trastuzumab experiment, patient’s tumor cells were resuspended in the antibody and the antibody was also added in the E3 medium (Table S2). At 1 dpi all zebrafish xenografts were screened regarding the presence of successfully injected tumor mass and randomly distributed in untreated-controls and treatment groups.

### Zebrafish-Xenograft treatment response evaluation

Apoptosis fold induction was quantified as the apoptosis fold change observed between treated and untreated zAvatars. Apoptosis percentage was calculated as follow:

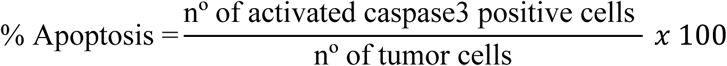

The apoptosis fold induction was calculated as follows:

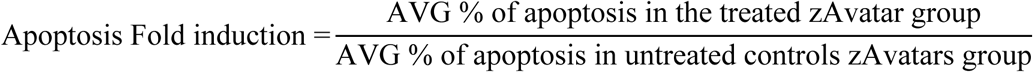

Apoptosis fold induction upon treatment was analyzed before knowing the patient’s clinical response. The zAvatar results were sent to the clinician to match results retrospectively. The selection of cut-off value for caspase3 Fold Induction was determined using the receiver operating characteristic (ROC) curve. This analysis was applied to assess the diagnostic performance of the zAvatars for differentiating patients into sensitive and resistant to the treatment. ROC analysis identified as an optimal threshold 1.4 apoptosis fold induction with highest specificity and sensitivity performance (AUC = 1), i.e., above, or equal to 1.4 of apoptosis fold induction we considered that the zAvatars were sensitive (S) to therapy, whereas below 1.4 were resistant (R) to the therapy. The area under the ROC curve (AUC) was calculated using XLSTAT software.

### Zebrafish xenograft whole-mount immunofluorescence

Primary antibodies: anti-Activated Caspase3 (rabbit, Cell signaling, 1:100, code#9661), anti-Human mitochondria (mouse, Merck Millipore, 1:100, cat#MAB1273), anti-Ki67 (mouse, Leica-Novocastra, 1:100, cat#NCL-Ki67-MM1), anti-GATA3 (mouse, SantaCruz, 1:100, cat#sc-269); anti-ER (mouse, SantaCruz; 1:100, cat#sc-8002); anti-PR (mouse, Santacruz, 1:100, cat#sc-166169), anti-Vimentin (mouse, Leica, 1:100, cat#PA0640); anti-FAP (mouse, Santacruz, 1:100, cat#65398); anti-α-SMA (mouse, Cell signaling, 1:100, cat#48938), anti-CK5 (rabbit, Abcam, 1:100, cat#ab52635), anti-CK18 (rabbit, Abcam, 1:100, cat#ab32118) anti-CD3 (mouse, Abcam, 1:100, cat#ab134096), anti-CD68 (mouse, Leica, ready-to-use, PA0273), and anti-CD163 (rabbit, Invitrogen, 1:100, PA5-78961). Secondary antibodies: Alexa Fluor 488 Phalloidin (Invitrogen, 1:400) Alexa goat anti-rabbit 488 (Molecular probes, 1:400), anti-mouse 488 (Molecular probes, 1:400), and anti-mouse 647 (Molecular probes 1:400) were applied simultaneously with DAPI (1:100). Xenografts were mounted with Mounting media.

### Imaging and quantification

All images were obtained in an upright Zeiss LSM 710 confocal microscope, generally with 5μm interval z stacks. Quantification analysis was performed using ImageJ software. Images were analyzed using ImageJ software, Cell counter plugin.^[55]^ Total number of cells / tumor = AVG (3 slices Zfirst, Zmiddle, Zlast) x total counted slices/1.5 for the Hs578T and MDA-MB-468 xenografts. In zAvatars, tumor cells were counted in each slice of the tumor. Mitosis and activated Caspase3 were quantified in all slices. zAvatars quantifications were performed in all tumor depth when tumor cells were spread, or the tumors were small. Whole zebrafish larvae images were composed using the stitching plugin from FIJI.^[56]^

### Metastatic potential quantification

The number of xenografts that had micrometastasis in the end of the assay was determined:

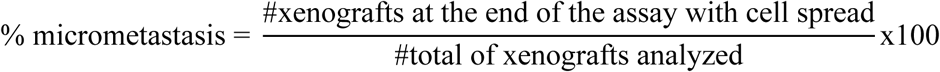

### Statistical analysis

Statistical analysis and graphical representation of the data were performed using GraphPad Prism 9.0 software. All data sets were challenged by a normality test (D’Agostino & Pearson and the Shapiro-Wilk). Data sets with a Gaussian distribution were analyzed by unpaired *t-*test. Datasets that did not pass the normality test were analyzed by Man-Whitney test. All normalized data (fold induction) were analyzed by Mann-Whitney test. Differences were considered statistically significant at P<0,05 and statistical outpour was represented by stars (*P<0.05, **P<0.01, ***P<0.001 and ****P< 0.0001) and non-significant (ns, P> 0,05). All data are presented as mean ± standard error of the mean (SEM) or standard deviation (SD).

### Institutional Review Board Statement

All experiments were approved by the Champalimaud Animal Ethical Committee and Portuguese institutional organizations—Órgão de Bem-Estar e Ética Animal/Animal Welfare and Ethics Body (ORBEA) and Direção Geral de Alimentação e Veterinária/Directorate General for Food and Veterinary (DGAV), following the 20130617 and 20200304 protocols.

### Informed Consent Statement

Informed consent was obtained from all subjects involved in the study.

## Statement

The data presented in this study are available on request from the corresponding author.

## Supporting Information

The data that supports the findings of this study are available in the Methods and Supporting Information of this article.

## Acknowledgements

We would like to thank all the patients who participated in the study. We would also like to thank all the Champalimaud Breast Unit, who approached patients for consent and collection of tissue as well as the Champalimaud Foundation Biobank (CFB). We acknowledge the Histopathology Unit of Champalimaud Clinical center and the Champalimaud Fish Platform. We thank M. Bettencourt-Dias Lab for the MDA-MB-468 and Hs578T cell lines.

## Author Contributions

R.F. conceptualized and supervised all research; M.G.F. co-supervised the initial research; R.M. and C.R.A. performed research; J.R., H.G. M.C-M. and M.J.B provided and analyzed patient’s clinical information; J.R., H.G. selected the patients for this study; M.C-M., M.J.B. and R.C-M. analyzed the histopathological features of the tumors; C.A., B.S., P.C., and M.J.C., provided tumor samples; E.B. provided and analyzed patient’s MRI and CT scans; F.C., H.G., J.R., M.C-M., M.J.B., R.C-M., C.A., B.S., P.C., and M.J.C., provided essential clinical advice and input throughout the study. R.M. and R.F. wrote the paper and all authors reviewed and approved it. Funding: R.F. and M.G.F.

## Data Availability Statement

The data that support the findings of this study are available upon author’s request.

## Supporting Information

### Supplementary Figures

**Figure S1.**
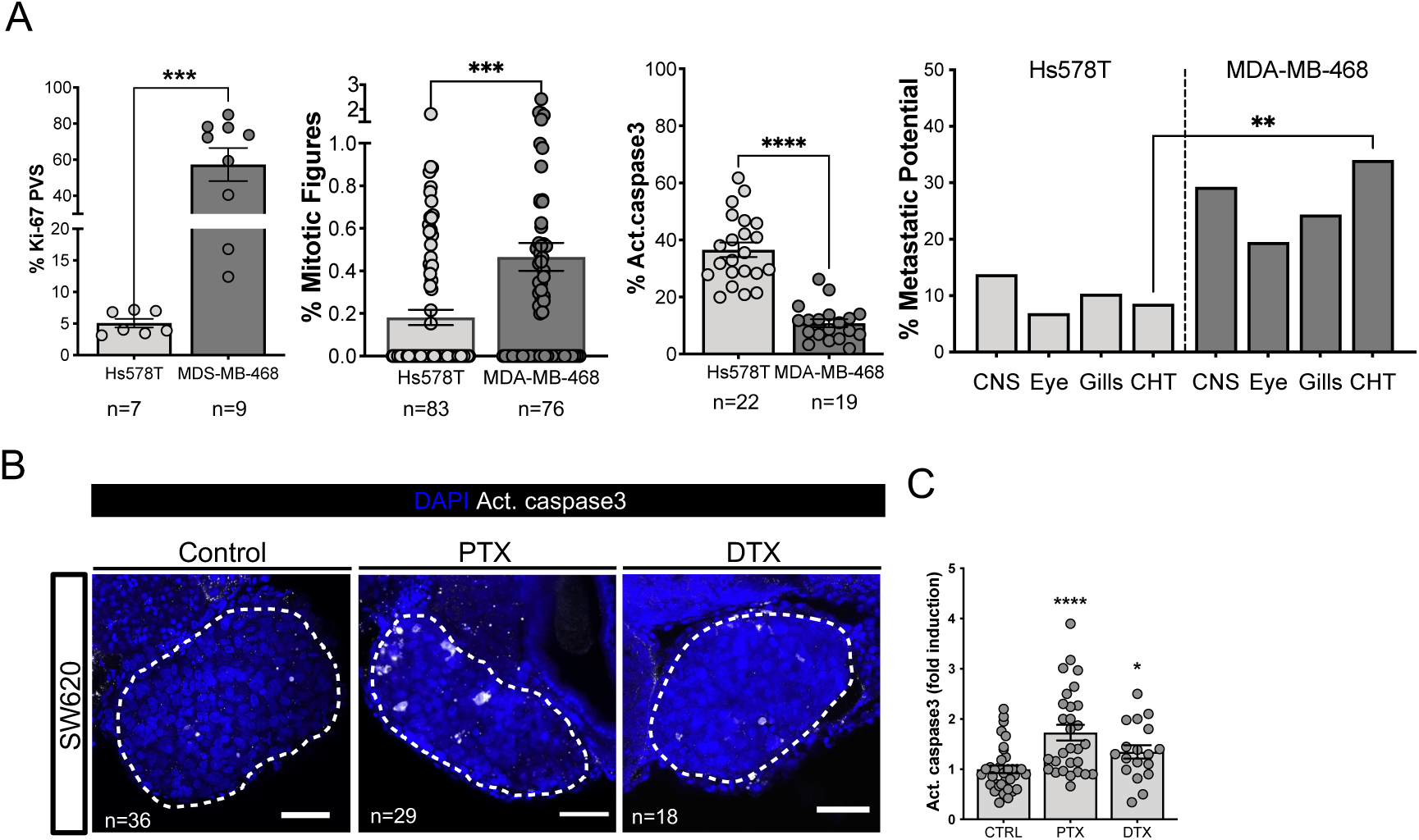
Characterization and comparison between two human TNBC tumor zebrafish xenografts and Colorectal cancer xenografts sensitivity to taxanes. A) Quantification in percentage of Ki-67 positive cells (AVG Hs578T = 5,1% vs AVG MDA-MB-468 = 57.6%), mitotic figures (AVG Hs578T = 0.18% vs AVG MDA-MB-468 = 0.46%) and activated caspase3 (AVG Hs578T = 36.57% vs AVG MDA-MB-468 = 10.9%) (each dot represents one xenograft). Quantification of the metastatic potential at 4dpi (AVG Hs578T = 8.6% vs AVG MDA-MB-468 = 34%) are expressed as AVG ± SEM. Hs578T (CNS n=29; Eye n=29; Gill n=29; CHT n=35), MDA-MB-468 (CNS n=41 Eye n=41; Gill n=41; CHT n=47). Results are expressed as AVG ± SEM and are averages from 1 and 2 (% activated caspase3) independent experiments. B), SW620 colorectal cancer zebrafish xenografts were treated in-vivo with paclitaxel or docetaxel and compared with non-treated controls. Zebrafish were fixed at 4 dpi, 3 days post-treatment (3dpt). C), Cell death by apoptosis (activated caspase3 in white) was analyzed, quantified, and normalized to the respective controls (*\*\*\*\*P<* 0.0001). Results are averages from three independent experiments, and the total number of xenografts analyzed is indicated in the images. Results are expressed as AVG±SEM are averages from one independent experiment. Each dot represents one xenograft. Statistical analysis was performed using Mann–Whitney test. White dashed line delimits the tumors. Statistical results \**P* < 0.05, \*\*\*\**P* < 0.0001. Scale bars represent 50 μm. All images are anterior to the left, posterior to right, dorsal up, and ventral down.

**Figure S2.**
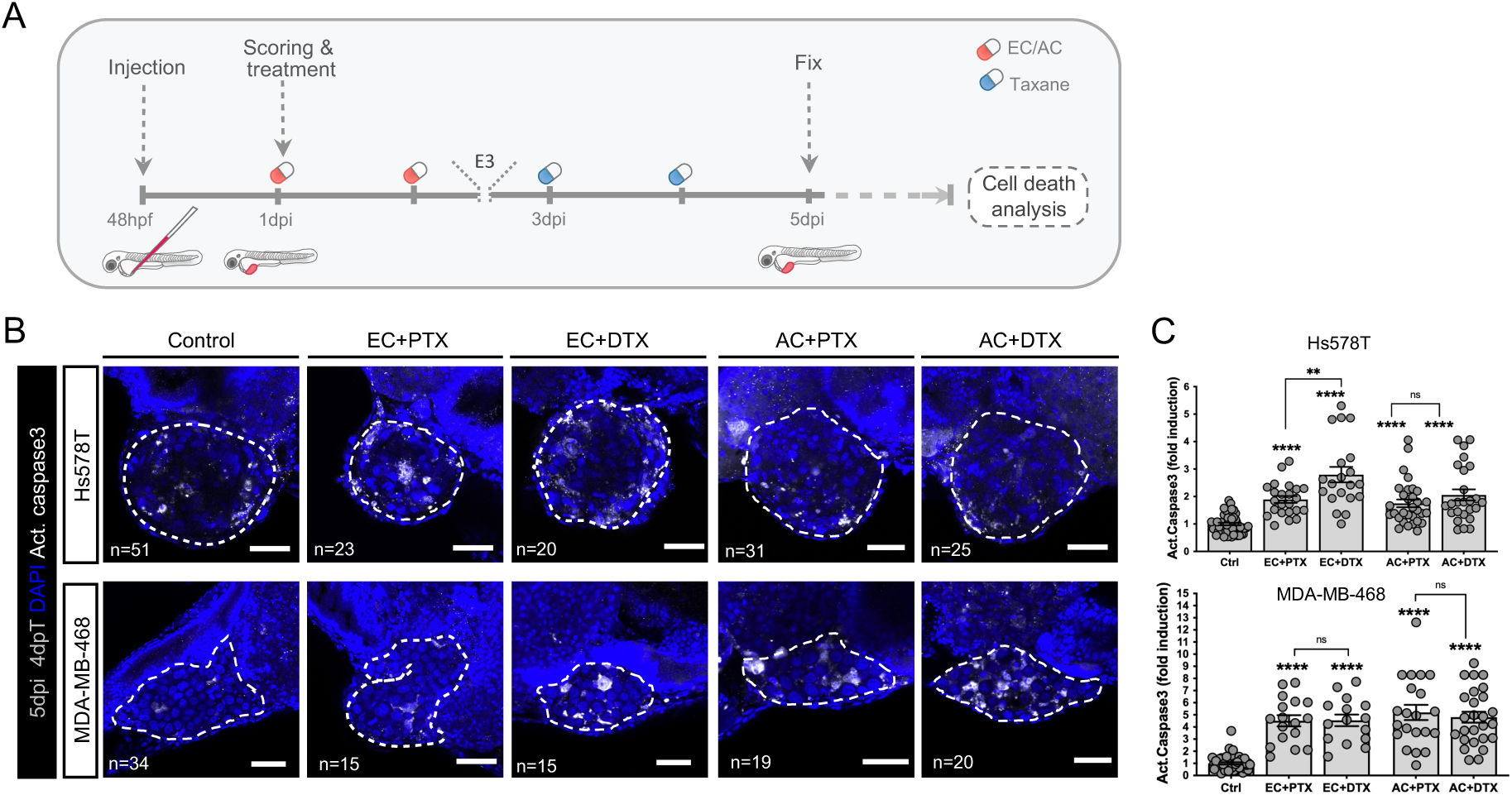
Single treatment with cyclophosphamide is as effective as its combination with anthracyclines and taxanes. A) Workflow depicting the therapy scheme: Patients’ samples were injected into zebrafish embryos at 48 hours post-fertilization (hpf), followed by tumor screening in the next day and treatment initiation. A treatment pause (E3) was introduced before switching chemotherapy regimens. Treatment was concluded at 5 days post-injection (dpi), with subsequent analysis of cell death using immunofluorescence. B) Human TNBC were injected in 48hpf zebrafish larvae, treated for 2 days with epirubicin + cyclophosphamide (EC) or doxorubicin + cyclophosphamide (AC), followed by a new round of 2 days with a taxane (paclitaxel or docetaxel) with a small pause in the treatments (E3) between EC/AC and taxanes. Xenografts were fixed at 5 dpi and processed for immunofluorescence. Hs578T and MDA-MB-468 breast cancer xenografts treated in-vivo with EC combined with either paclitaxel or docetaxel, or AC combined with either paclitaxel or docetaxel. C), Cell death by apoptosis (activated caspase3 in white) was analyzed and quantified (*\*\*P=*0.0073, \*\*\*\**P* < 0.0001). Results are expressed as AVG±SEM and are from three independent experiments. Each dot represents one xenograft. The total number of xenografts analyzed is indicated in the images. White dashed line delimits the tumors. Statistical analysis was performed using Mann–Whitney test. Statistical results: (ns) > 0.05, \**P* < 0.05, \*\**P* < 0.01, \*\*\**P* < 0.001, \*\*\*\**P* < 0.0001. Scale bars represent 50 μm. E3: Embryo medium. All images are anterior to the left, posterior to right, dorsal up, and ventral down.

**Figure S3.**
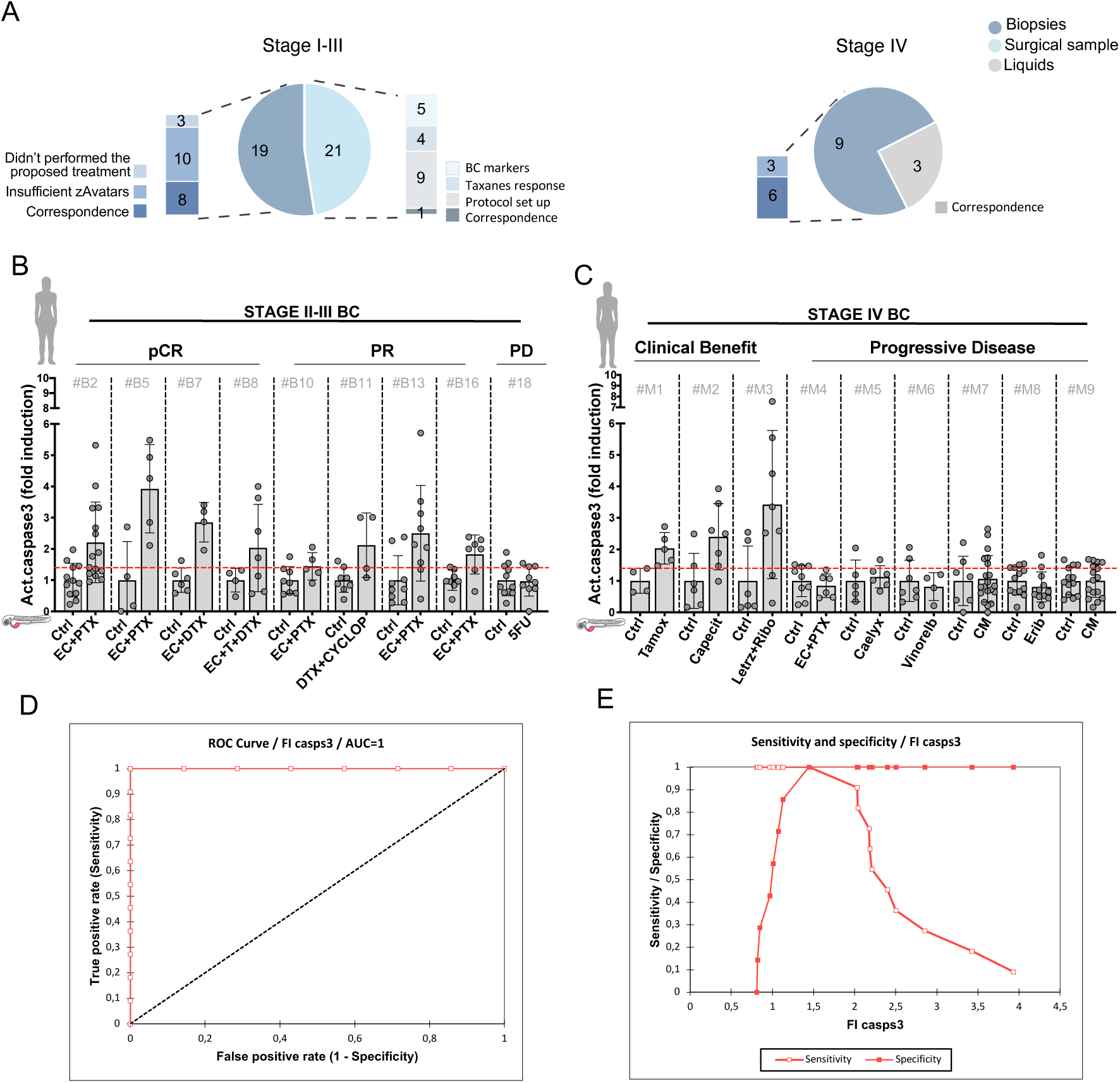
zAvatars response to therapy can predict patient’s clinical response. A), Pie charts represent all BC samples injected into the zebrafish embryos. B), 2dpf zebrafish were injected with core-needle breast cancer biopsies from 8 patients with early BC that underwent NA chemotherapy: 4 (#B2, #B5, #B7 and #B8) achieved a pCR, the other 4 patients achieve a PR (#B10, #B11, #B13 and #B16) and 1 patient from surgical sample showed PD (#18). Cell death by apoptosis (activated caspase3) was analyzed. C), 2dpf zebrafish were injected with samples from advanced (stage IV) BC: 3 achieved CB (#M1 - #M3) and the remaining 6 had PD (#M4 - #M9). D), Receiver operating characteristics curve (ROC) analysis of similarity between zPDX response and patients’ response to the treatment AUC is 1 (95% CI) for activated caspase3 fold induction. (E), Graph result showing Sensitivity and Specificity / fold induction caspase3. #B2 - #B16: core needle biopsies from breast; #18: surgical samples; #M1, #M3 and #M4– core needle biopsies from breast; #M2 and #M6 – core needle biopsy from liver; #M5 – core needle biopsies from lymph node; #M7 – Ascites, and #M8 and #M9 – Pleural effusions. pCR - pathologic complete response; PR - partial response; PD - progressive disease; CB - clinical benefit; PD – progressive disease.

**Figure S4.**
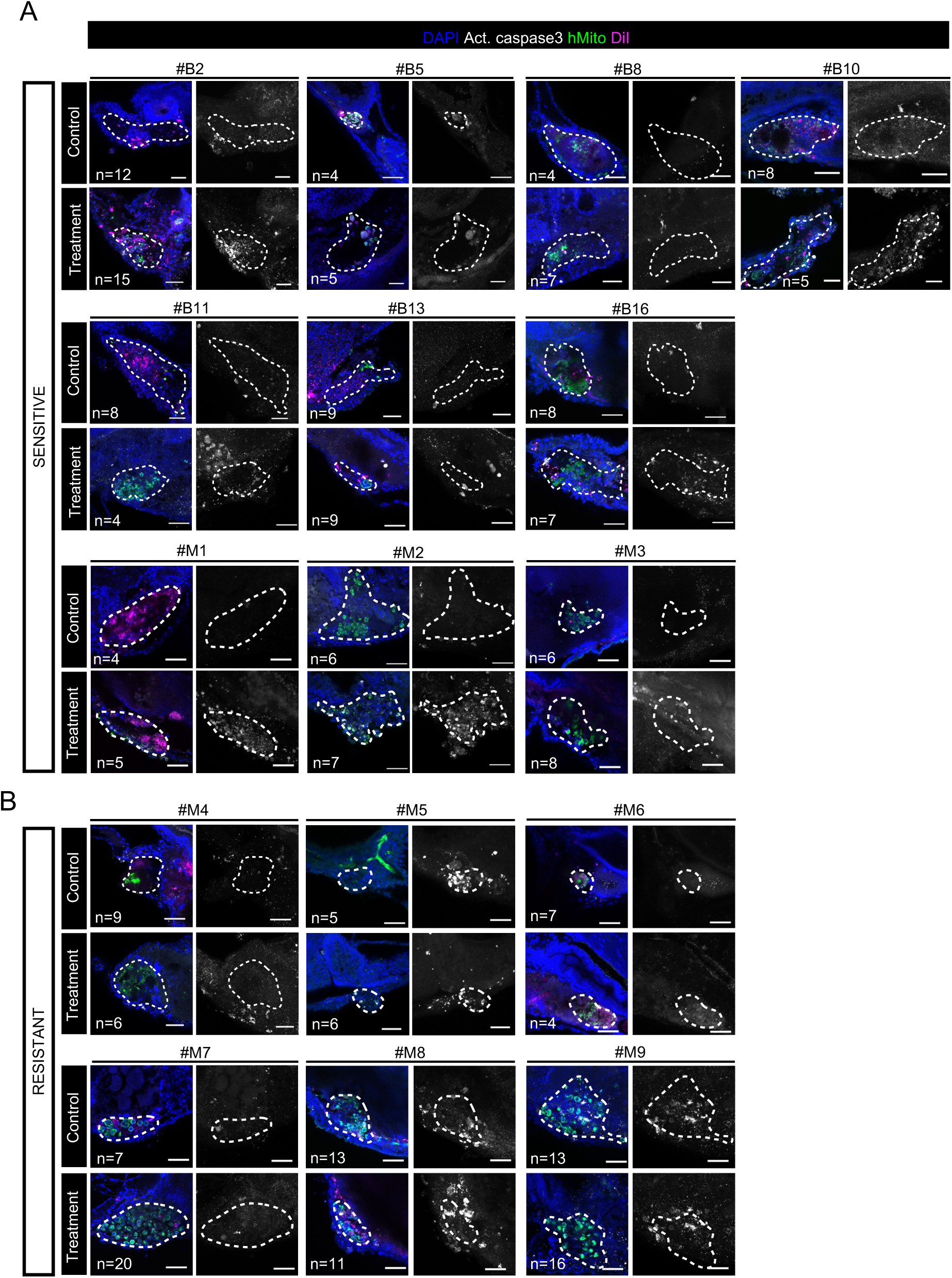
zAvatars response to therapy can predict patient’s clinical response. A), Sensitive BC zAvatars from early stage: primary breast core needle-biopsies (#B2 - #B16); advanced stage: breast core needle-biopsy (#M1 and #M3), liver core needle-biopsy (#M2). zAvatars treated in-vivo with the same ChT regimen as its donor-patient and compared with non-treated controls. Zebrafish were fixed at 3, 4 or 5dpi, corresponding to 2, 3 or 4days post-treatment (dpt). B), Resistant metastatic BC zAvatars from stage IV: breast core needle-biopsy (#M4), lymph-node core needle-biopsy (#M5), liver core needle-biopsy (#M6), drainage from ascites (#M7) and MPE (#M8 and #M9). zAvatars treated in-vivo with the same ChT regimen as its donor-patient and compared with untreated controls. zAvatars were fixed at 3 or 4dpi, corresponding to 2- or 3-days post-treatment (dpt). dpt- days post-treatment. MPE-malignant pleural effusion. Cell death by apoptosis (activated caspase3 in white, Z-projection image). Total number of xenografts analyzed are indicated in the images. White dashed line delimits the tumors. Scale bars represent 50 μm. All images are anterior to the left, posterior to right, dorsal up, and ventral down.

**Figure S5.**
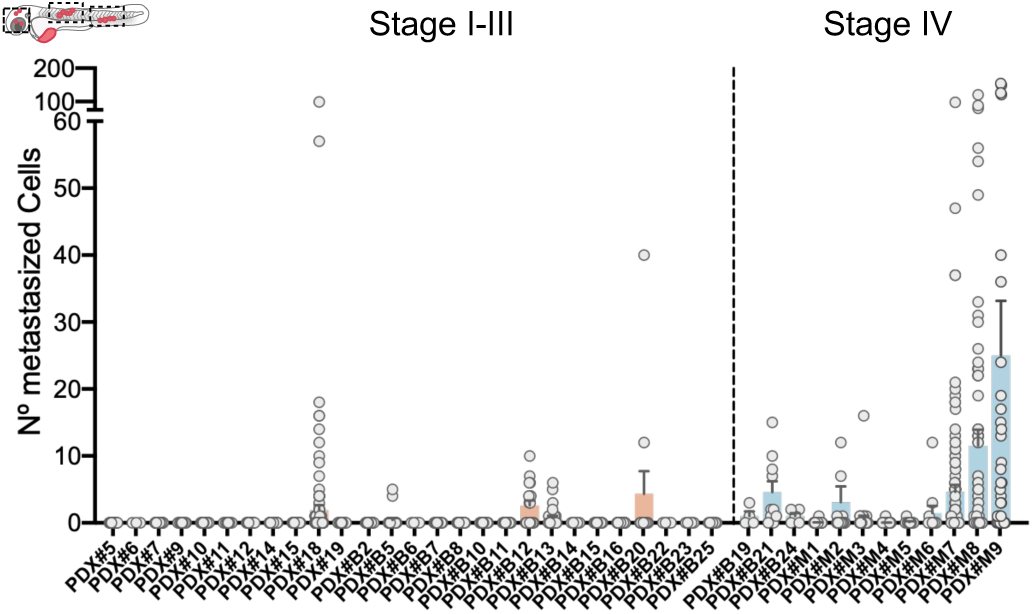
Micrometastasis dissemination in zAvatars from stage I-IV breast cancer. A), Quantification of the number of tumor cells disseminated throughout the zAvatar at 3, 4 or 5dpi without treatment (controls) generated from patients staged (I-IV). zAvatars were analyzed in a fluorescent stereoscope to detect micrometastasis. Each dot represents the number of micrometastasis in the zAvatars. The outcomes are expressed as AVG ± SEM. Number of zAvatars per patient ranged between 4 - 50. Data from one independent experiment.

**Table S1.**
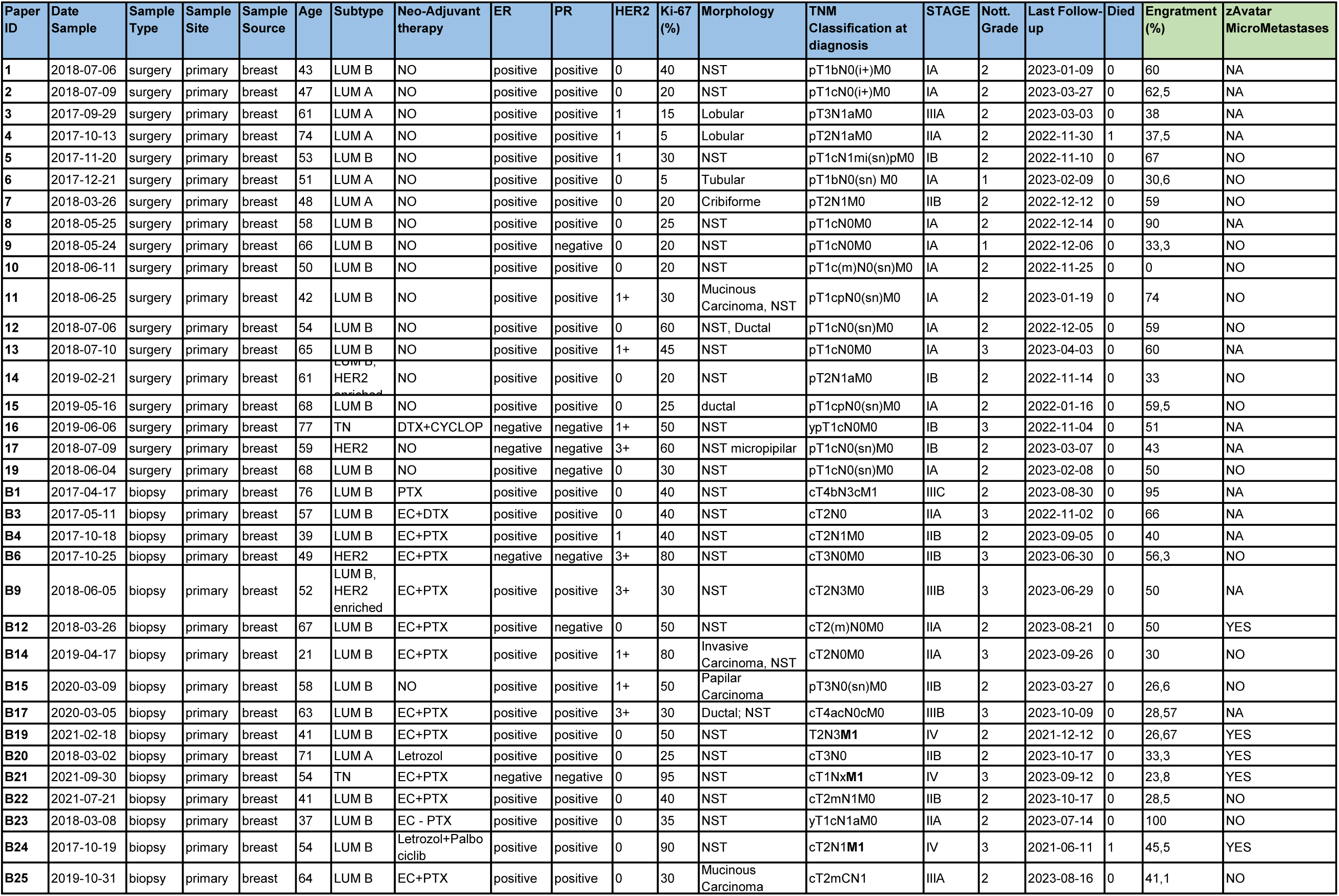
Patient’s Clinical Information and correcponding zAvatars-test results.

**Table S2.**
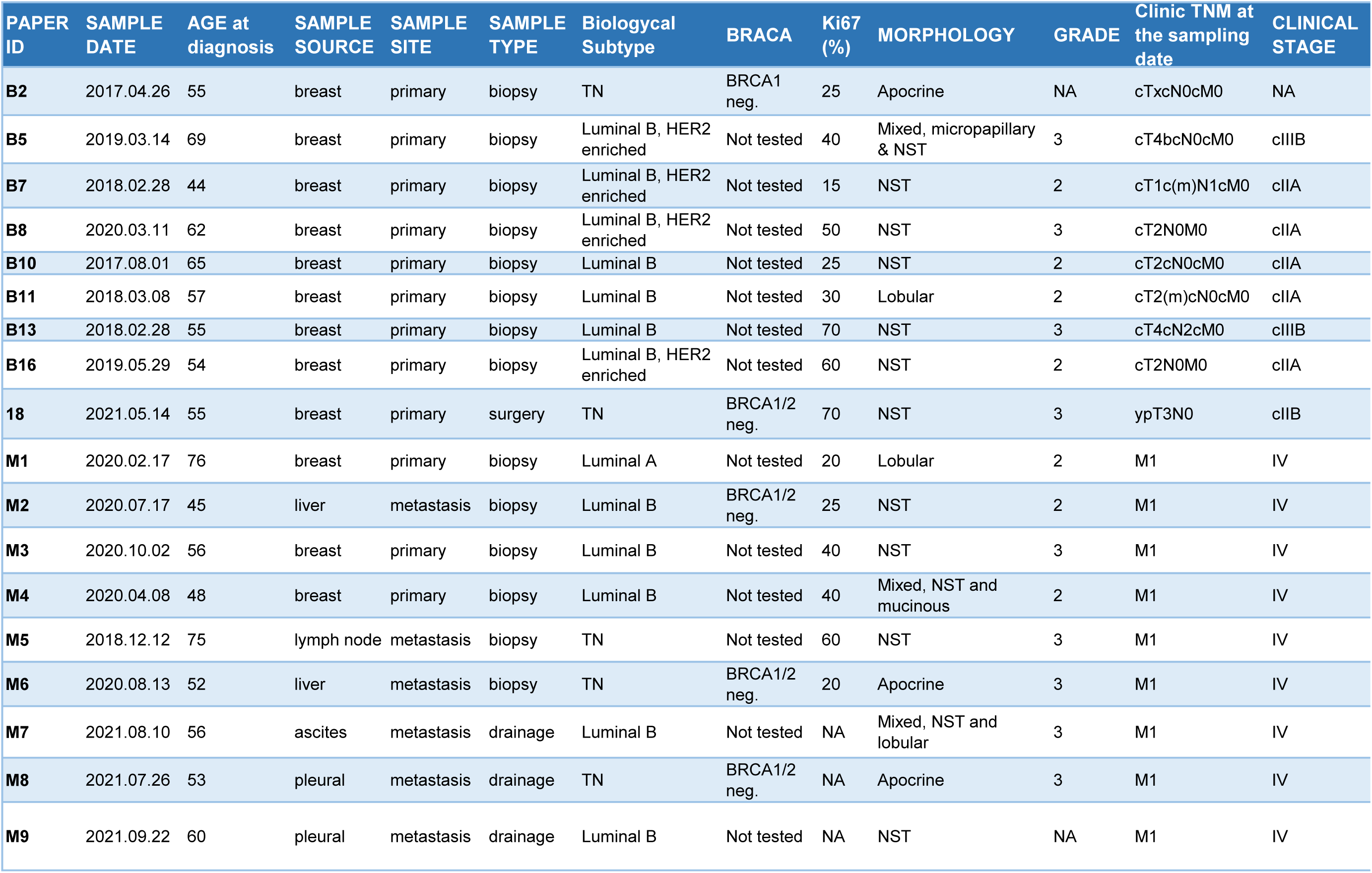
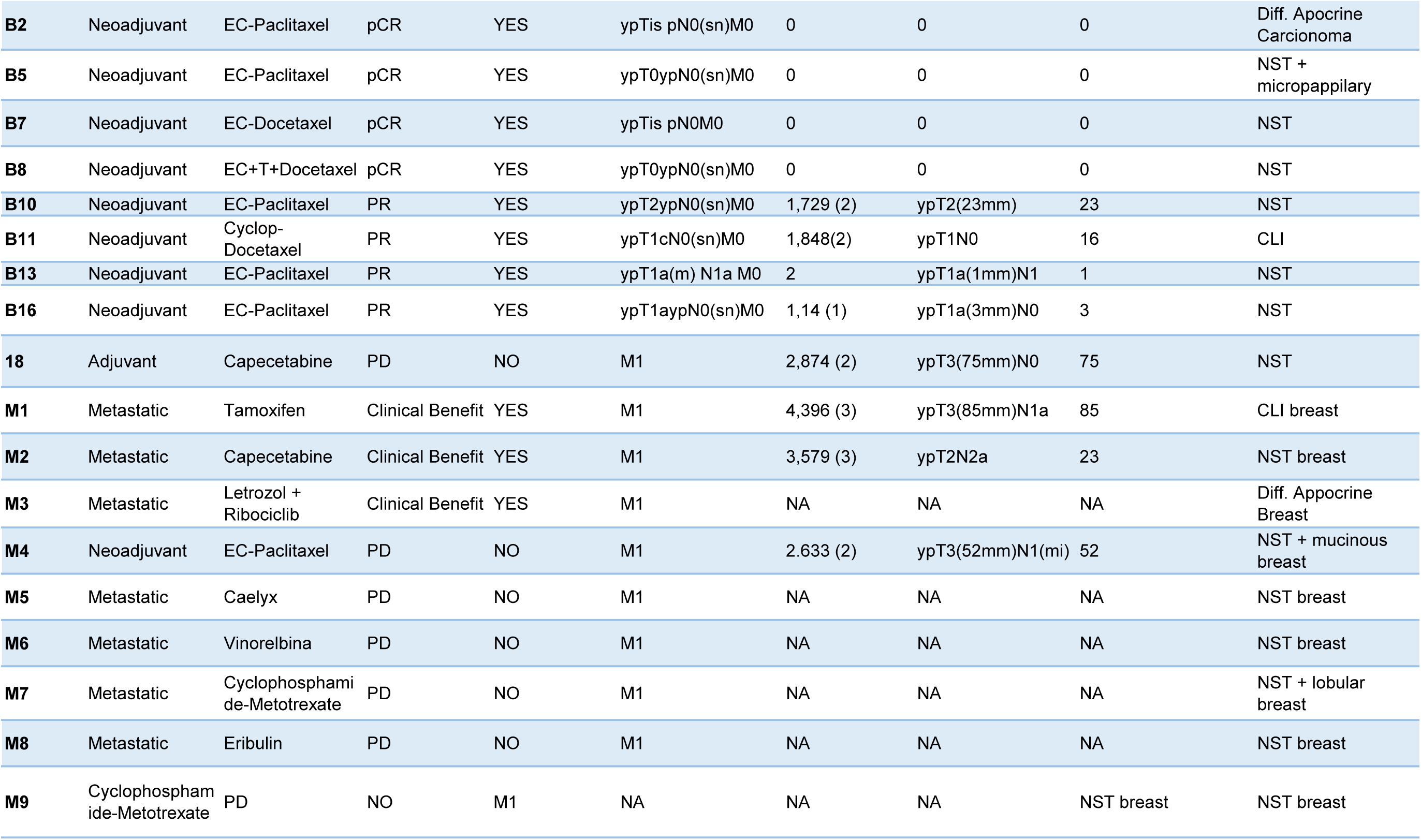
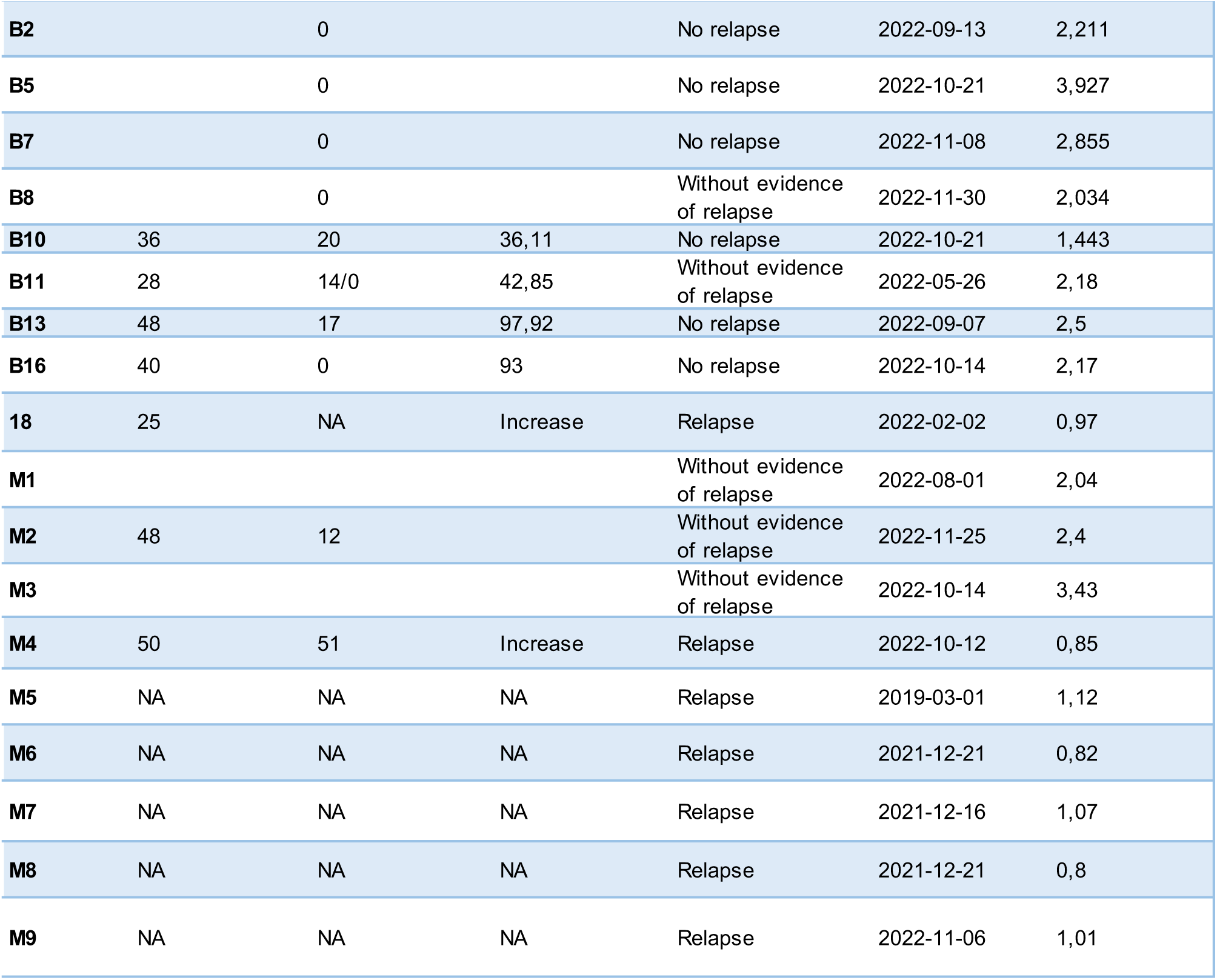
Clinical Information of Patients used for zAvatar correspondence.

**Table S3.** Chemotherapies used for zAvatars treatment.

|  | **Chemotherapy scheme** | **Drug** | **Concentration** | **REF** |
| --- | --- | --- | --- | --- |
| **Cell derived xenografts** | Paclitaxel (PTX) |  | 140mM | [57] |
|  | Docetaxel (DTX) | - | 46,8mM | [58] |
|  | Epirubicin | - | 68,9mM | [59] |
|  | Doxorubicin | - | 73,56uM | [59] |
|  | Cyclophosphamide | - | 1,43mM | [60] |
|  | Epirubicin + Cyclophosphamide (EC) | Epirubicin | 13,8mM + | [59] |
|  |  | Cyclophosphamide | 280uM | [60] |
|  | Doxorubicin + Cyclophosphamide (AC) | Doxorubicin | 15uM + | [59] |
|  |  | Cyclophosphamide | 280uM | [60] |
|  | EC + PTX | - | 13,8mM + 280uM + 70mM | [59], [60], [57] |
|  | EC + DTX | - | 13,8mM + 280uM + 19mM | [59], [60], [58] |
|  | AC + PTX | - | 15uM + 280uM + 70mM | [58], [59] |
|  | AC + DTX | - | 15uM + 280uM + 19mM | [61], [58] |
| **Patient derived xenografts** | Anthracycline + Cyclophosphamide – Taxane (DTX / PTX) + Trastuzumab | Anthracycline (Epirubicin) | 13,8mM | [59] |
|  |  | Cyclophosphamide | 280uM | [60] |
|  |  | DTX / PTX | 15,4mM / 70mM | [58]/[57] |
|  |  | Trastuzumab | 15µg·mL<sup>-1</sup> (cells)<br>250µg·mL<sup>-1</sup> (E3) | [62] |
|  | CM (Cyclophosphamide + Methotrexate) | Cyclophosphamide | 280uM | [60] |
|  |  | Methotrexate | 7,5mM | [63] |
|  | Anthracycline + Cyclophosphamide – Taxane (DTX / PTX) | Anthracycline (Epirubicin) | 13,8mM | [59] |
|  |  | Cyclophosphamide | 280uM | [60] |
|  |  | DTX / PTX | 15,4mM / 70mM | [58]/[57] |
|  | Docetaxel + Cyclophosphamide | DTX | 15,4mM | [58] |
|  |  | Cyclophosphamide | 280uM | [60] |
|  | Tamoxifen | Tamoxifen | 10uM | [64] |
|  | 5FU | 5FU | 4,2mM | [65] |
|  | Letrozol + Ribociclib | Letrozol | 911nM | [66] |
|  |  | Ribociclib | 150uM | [67] |
|  | Caelyx (Doxor) | Caelyx | 3mM | [61] |
|  | Vinorelbine | Vinorelbine | 186uM | [68] |
|  | Eribulin | Eribulin | 4mM | [69] |

**Table S4.**
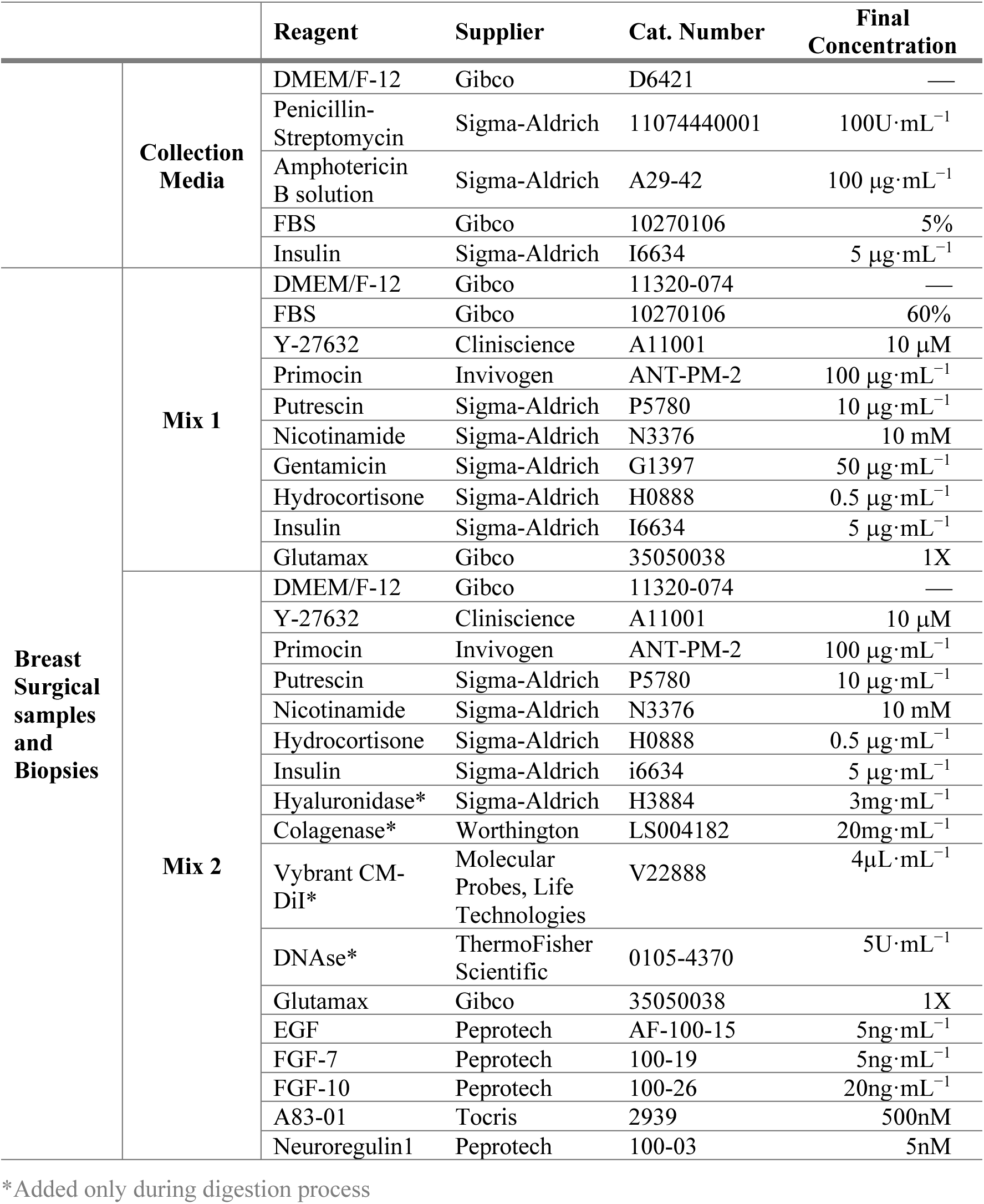
Compounds used for zAvatars generation.

**Table S5.** Compounds used for zAvatars generation.

|  |  | **Reagent** | **Supplier** | **Cat. Number** | **Final Concentration** |
| --- | --- | --- | --- | --- | --- |
| **Malignant pleural effusion and Ascites** | Mix 1 | DMEM/F-12 | Gibco | 11320-074 | — |
|  |  | B27 | Gibco | 17504-044 | 1X |
| | | Y-27632 | Cliniscience | A11001 | 10μM |
| | | Primocin | Invivogen | ANT-PM-2 | 100μg·mL<sup>-1</sup> |
| | | Putrescin | Sigma-Aldrich | P5780 | 10μg·mL<sup>-1</sup> |
|  |  | Nicotinamide | Sigma-Aldrich | N3376 | 10mM |
| | | Hydrocortisone | Sigma-Aldrich | H0888 | 0.5μg·mL<sup>-1</sup> |
| | | Insulin | Sigma-Aldrich | I6634 | 5μg·mL<sup>-1</sup> |
|  |  | Glutamax | Gibco | 35050038 | 1X |
|  |  | Benzonase | ThermoFisher Scientific | 88700 | 37.5U/ml |
|  |  | NEAA | ThermoFisher Scientific | 11140050 | 1X |
| | | Vybrant CM-DiI* | Molecular Probes, Life Technologies | V22888 | 4μL·mL<sup>-1</sup> |
| | | Noggin | Peprotech | 120-10C | 0.2μg·mL<sup>-1</sup> |
|  |  | NAC | Sigma-Aldrich | A0737 | 1.25mM |
| | | Heregulinβ-1 | Peprotech | 100-03 | 5nM |
|  |  | FGF-10 | Peprotech | 100-26 | 20ng·mL<sup>-1</sup> |
|  |  | FGF-7 | Peprotech | 100-19 | 5ng·mL<sup>-1</sup> |
|  |  | EGF | Peprotech | 100-15 | 5ng·mL<sup>-1</sup> |
|  |  | A83-01 | Tocris | 2939 | 500nM |

\*Added only during digestion process

